# Pectin digestion in herbivorous beetles: Impact of pseudoenzymes exceeds that of their active counterparts

**DOI:** 10.1101/462531

**Authors:** Roy Kirsch, Grit Kunert, Heiko Vogel, Yannick Pauchet

**Affiliations:** Department of Entomology Max Planck Institute for Chemical Ecology, Hans-Knoell-Str. 8, Jena, 07745, Germany; Department of Biochemistry Max Planck Institute for Chemical Ecology, Hans-Knoell-Str. 8, Jena, 07745, Germany

## Abstract

Many protein families harbor pseudoenzymes that have lost the catalytic function of their enzymatically active counterparts. Assigning alternative function and importance to these proteins is challenging [1]. Because the evolution towards pseudoenzymes is driven by gene duplication, they often accumulate in multigene families. Plant cell wall-degrading enzymes (PCWDEs) are prominent examples of expanded gene families. The pectolytic glycoside hydrolase family 28 (GH28) allows herbivorous insects to break down the PCW polysaccharide pectin. GH28 in the Phytophaga clade of beetles contains many active enzymes but also many inactive counterparts. Using functional characterization, gene silencing, global transcriptome analyses and recordings of life history traits, we found that not only catalytically active but also inactive GH28 proteins are part of the same pectin-digesting pathway. The robustness and plasticity of this pathway and thus its importance for the beetle is supported by extremely high steady-state expression levels and counter-regulatory mechanisms. Unexpectedly, the impact of pseudoenzymes on the pectin-digesting pathway in Phytophaga beetles exceeds even the influence of their active counterparts, such as a lowered efficiency of food-to-energy conversion and a prolongation of the developmental period.

## Introduction

Though plants contain all the nutrients herbivorous insects need, dependence on plants as a food source is challenging for two reasons: nutrient amounts and ratios are highly variable, and nutrient requirements are not uniform over an insect’s life cycle [2]. Because a large proportion of ingested food consists of macromolecules -- proteins and polysaccharides -- which can be limited in a plant diet, herbivorous insects have adapted to this scarcity by evolving specific digestive capacities [3-5]. The intake of these macromolecules is regulated [6] before their degradation by hydrolases to release amino acids and sugars that can be absorbed and used by insects as sources of nitrogen and metabolic energy, respectively. Starch, the main storage polysaccharide in plants, is of great importance as the source of energy, which fuels insect growth and development. This polysaccharide can be easily digested by amylases, which are widespread in herbivorous insects [7]. In addition to starch, plant cell wall (PCW) polysaccharides are major carbohydrate constituents of green plants and can make up half of a leaf’s dry weight [8]. Every growing plant cell is encased in a primary wall that is composed of approximately 90 % polysaccharides [9]. PCW polysaccharides include cellulose and hemicellulose fibers that are further embedded in a pectin polysaccharide matrix [10]. This carbohydrate network can be broken down by plant cell wall-degrading enzymes (PCWDEs), which mainly belong to different families of carbohydrate esterases, polysaccharide lyases and glycoside hydrolases [11]. For a long time, indirect evidence related to the activity of insects’ endogenous PCWDE has accumulated, but despite more recent attempts to clone and characterize these enzymes [12], little is known about their relevance in herbivorous insects with respect to nutrient acquisition.

In contrast to the more limited information available for their role in insects, PCWDEs have been extensively studied in plant-pathogenic microbes [13, 14], in particular, pectin-degrading polygalacturonases (PGs), which belong to the glycoside hydrolase family 28 (GH28). Genes encoding GH28 PGs are widespread in plant pathogens [15], and they were present in the most recent common ancestor of fungi, which illustrates their primary importance at the pathogen-plant interface [16]. PGs have been shown to be key players during plant infestation by these microbes, as (i) they are the first enzymes secreted to weaken the plant cell wall and (ii) they are important virulence factors [17-20].

More recently, endogenous - and apparently widespread – PG encoding genes were identified in a number of herbivorous insect orders, including Hemiptera (mirid bugs), Phasmatodea (stick insects), Hymenoptera (gall wasps) and Coleoptera (mainly “Phytophaga”: see below) [21-27]. Functional characterization *in vitro* revealed that the corresponding proteins were active PGs that hydrolyze the homogalacturonan pectin backbone synergistically, releasing galacturonic acid oligomers and monomers [24, 25, 27, 28]. However in beetles, inactive GH28s were identified that cannot bind their predicted pectin substrate due to amino acid substitutions at crucial positions [24, 28]. Despite this, all GH28 family members of beetles and stick insects were shown to be specifically expressed in gut tissue, with the corresponding enzymes being secreted into the gut lumen [28-32]. PGs of mirid bugs are expressed in salivary glands and are injected into the plant during piercing [22, 33]. Taken together, these factors strongly indicate a digestive function for PGs and their central role in pectin hydrolysis *in vivo*.

Remarkably, insect genes encoding PGs seemed to be acquired by horizontal gene transfer (HGT) [24, 27, 28]. Especially in the “Phytophaga” beetles -- the hyper-diverse beetle clade that includes weevils, longhorned beetles and leaf beetles [34] -- several rounds of loss and replacement have affected the evolutionary history of the beetle GH28 gene family [24]. PGs persisted after the initial HGT early in the evolution of “Phytophaga” beetles, and their genes have duplicated and are under continued action of purifying selection while the corresponding proteins have functionally diversified. These facts strongly indicate that an important function of PGs is to promote herbivory in this clade of beetles, which represents about 50 % of all herbivorous insects [35].

Additionally, symbionts of herbivorous insects can encode for pectinase activity [12, 36]. A remarkable example shows how a “Phytophaga” beetle host benefits from a symbiont’s pectolytic activity [37]: when the pectinase-encoding symbiont is removed from its *Cassida rubiginosa* leaf beetle host, insect survival is highly reduced. The symbiont most likely compensates for the loss of the *Cassida* endogenous PG genes, and host survival depends on pectin digestion facilitated by the symbiont. The lack of beetle PGs in the Phytophaga clade, as in *Cassida*, is so far unusual. However, this system clearly exemplifies the importance of a single PCWDE family for the fitness of *C. rubiginosa* beetles.

To directly test the biological relevance of the endogenous pectin-degrading ability of an insect, we analyze the effects of gene silencing on (i) the performance of a leaf beetle, (ii) the enzymatic activities and (iii) the global gene expression. We simultaneously silenced the three endo-PGs of the mustard leaf beetle *Phaedon cochleariae*, and, in another RNAi treatment, three GH28 family members that had lost their PG enzymatic activity [24]. This simultaneous knock down, enabled us to test whether inactive GH28 family members continue to play a role in pectin hydrolysis, their ancestral function, even if they are not hydrolyzing polygalacturonan. Gene silencing allows us to study the significance of active enzymes and their pseudoenzyme counterparts and to understand why they become inactive towards a substrate during evolution while still under strong purifying selection [24].

## Material & Methods

### Insect and plant rearing

*Phaedon cochleariae* was reared in the laboratory (15 °C, long day conditions, 16-h/8-h light/dark period) on Chinese cabbage (*Brassica rapa* ssp. *pekinensis*) leaves for several generations. Larvae used for RNA interference (RNAi) experiments stemmed from an over-night egg laying of mass-reared adults. Egg-containing leaves were separated, and emerging larvae were fed with Chinese cabbage until injected with double-stranded RNA. Cabbage plants used for bioassays (*B. rapa* ssp. *pekinensis* var. Cantonner Witkropp) were reared in the greenhouse (21 °C, 55 % humidity, long day conditions, 14-h/10-h light/dark period), and larvae were fed with middle-aged leaves from 6- to 8-week-old non-flowering plants.

### Heterologous expression and enzymatic assays

Sf9 insect cells were cultivated in GIBCO Sf-900 II SFM (Invitrogen) on 6-well plates at 27 °C until 70-90% confluence was achieved. Transfection was performed with FuGENE^®^ HD (Promega) following the manufacturer’s protocol using the GH28 pIB/V5-His TOPO TA (Thermo Scientific) constructs described previously [24]. At 72 h after transfection, the culture medium of Sf9 cells was harvested and concentrated 10-fold using Pierce Concentrators 20 ml with a 20 kDa cutoff (Thermo Fisher Scientific, US). Culture medium was further dialyzed against water at 4 °C for 48 h using Slide-A-Lyzer Dialysis Cassettes with a 10 kDa cutoff, followed by desalting with Zeba Desalt Spin Columns with a 7 kDa cutoff (both Thermo Fisher Scientific) according to the manufacturer’s guidelines. The crude protein extract of transient heterologously expressed GH28 family members was used for Western blot analyses and enzyme assays. Expressed proteins were detected by Western blots using an anti-V5 HRP antibody (Thermo Fisher Scientific) and the SuperSignal West HisProbe Kit (Pierce, Germany).

For qualitative analysis of breakdown products of the beetle GH28 family members by thin layer chromatography (TLC), 16 μl of a 2.5% (w/v) PCW suspension was added to 20 μl GH28 and 4 μl citrate-phosphate buffer pH 5.0 to a final concentration of 20 mM. In the negative control (-), GH28 was substituted with 20 μl of water. Assays were analyzed as previously described [24]. Chinese cabbage PCW substrate was prepared after PCW enrichment and protein extraction as described for *Arabidopsis* hypocotyls [38]. After the protein supernatant was separated from the PCW pellet, the lyophilized and re-suspended pellet was used for enzymatic assays.

### RNA interference

Double-stranded RNAs of PCO-GH28-1, 3, 5, 6, 7 and 9 were prepared using the MEGAscript RNAi Kit (Life Technologies, Germany) according to the manufacturer’s instructions. Templates for synthesis were amplified from the corresponding expression vectors (pIBV5) from a previous study [24] using primers with overhangs containing the minimum T7 polymerase promotor sequence needed for transcription (Table S1). Off-target effects were predicted by searching all possible 21-mers of both RNA strands against our in-house *P*. *cochleariae* transcriptome database, allowing for one mismatch. Five-day-old larvae (early 2nd instar) were weighed before injection, and only those weighing 1.1-1.4 mg were used to ensure high survivorship (determines the lower limit) and at the same time a maximum of days in the larval stage (determines the upper limit). Larvae were immobilized on sticky tape and injected with 100 nL containing 100 ng of dsRNA of each GH28 as a pool or 300 ng of dsRNA of GFP using a Nano2010 injector (World Precision Instruments, US) oriented with a manual micromanipulator. Injected and non-injected larvae were kept in clear, ventilated plastic boxes (20×20×6 cm) containing a moistened tissue and cabbage leaves as food under standard rearing conditions.

### Monitoring larval development and food consumption

One day post injection, larvae were weighed and transferred individually (n=50 for each treatment: GH28-active (28a), GH28-inactive (28i), GFP injection control) to Petri dishes (diameter: 60mm) in order to document food consumption and development. Petri dishes, equipped with a leaf disc (diameter: 18mm (day 1-4 pi), 20mm (day 4-6 pi), 22mm (day 6-7 post injection)) on a filter paper moistened with 100μl sterile water, were sealed with Parafilm M to prevent desiccation and kept under standard rearing conditions. Leaf discs were changed every day and food was available *ad libitum*. Larval weight was recorded on days 1 and 5 post injection to calculate larval weight gain. Leaf discs were photographed every day to calculate food consumption (cm2 leaf area) by image analysis [39].

### Sample preparation

Five days post injection, larvae were dissected for gene expression analysis and enzymatic assays. Dissection was executed in 20 mM citrate/phosphate buffer pH 5.0 containing a cocktail of protease inhibitors (Complete EDTA-free, Roche, Germany). Intact whole guts were transferred in 200 μl of the same buffer chilled on ice, opened on one side and soaked in the buffer. The resulting buffer/gut content mixture was kept on ice during gut dissection and immediately centrifuged afterwards (5,000 g, 5 min, 4 °C). The supernatant was collected and stored at −20 °C until use. The remaining gut tissue was transferred to 450 μl RL buffer of the RNA extraction kit and frozen at -20 °C. Five replicates per treatment of four larval gut tissues and gut content each were taken for downstream analyses of gene expression and PG activity.

### Expression analyses

To compare gene expression in larvae injected with dsRNA targeting the active and inactive GH28 family members with GFP control, real-time quantitative PCR was performed. Each assay was set up in two technical replicates for each of the five biological replicates. As *P*. *cochleariae* has nine PG family members [24], we included both the silenced and the non-silenced ones in our analyses. RNA extraction was performed using the innuPrep DNA/RNA Mini kit (analytikjena) following the manufacturer’s instructions. After RNA integrity on a 1% agarose gel was checked, 500 ng of total RNA from each pool was reverse-transcribed with a 3:1 mix of random and oligo-dT20 primers. RT-qPCR was performed in optical 96-well plates on a CFX Connect detection system (BioRad, US). All steps were performed with the Verso SYBR Green 2-Step QRT-PCR Kit (Thermo Fisher Scientific, US) following the manufacturer’s instructions. The PCR program was as follows: 95 °C for 15 min, then 40 cycles at 95 °C for 15 s, 58 °C for 30 s, and 72 °C for 30 s, and afterward a melt cycle from 55 to 95 °C in 0.5-s increments. All primers were designed using Primer3 (version 0.4.0) and are listed in Supplementary Table S1. Specific amplification of each transcript was verified by dissociation curve analysis. A standard curve for each primer pair was determined in the CFX Manager (version 3.1) based on Cq values (quantitation cycle) of qPCR running with a dilution series of cDNA pools. The efficiency and amplification factors of each qPCR primer pair based on the slope of the standard curve was calculated with the help of the efficiency calculator (http://www.thermoscientificbio.com/webtools/qpcrefficiency/). *Elongation factor 1α* (*EF1α*; HE962191) was used as reference gene, and quantities of the genes of interest were expressed as RNA molecules of GOI/1000 RNA molecules of EF1α. The Cq values were determined from two technical replicates of each of the five biological replicates, and error bars indicate the standard error of the mean.

### Quantification of total polygalacturonase activity of gut content

Gut content samples were desalted with Zeba Desalt Spin Columns with a 7 kDa cutoff (Thermo Fisher Scientific, US) according to the manufacturer’s guidelines, and protein concentrations were determined using the Bradford reagent [40]. Quantitative assays measuring the release of reducing sugars after the hydrolysis of polygalacturonic acid were set up and analyzed as previously described (Kirsch et al., 2016) with slight modifications. In detail, 500 ng gut content protein was incubated with 0.2% polygalacturonic acid in a 20 mM citrate/phosphate buffer pH 5.0 at 40 °C for 10, 30, 60, 120 and 240 min. As a negative control, protein samples were boiled before incubation. Each assay was set up in three technical replicates for each of the five biological replicates, and reducing groups released after substrate hydrolysis were quantified by the 3,5-dinitrosalicylic acid (DNS) method [41].

Three solutions were prepared for analysis of these samples in advance as follows: solution 1 containing DNS, phenol and sodium hydroxide to a final concentration of 1%, 0.2% and 1% (w/v) in water, respectively. Solution 2 is a 100-fold stock of sodium sulfite to a final concentration of 0.5% (w/v) in water, and solution 3 is a 7-fold stock of potassium sodium tartrate (Rochelle Salt) to a final concentration of 40% (w/v) in water. A mixture of solutions 1 and 2 in a 99:1 ratio (v/v) was prepared fresh each time just before use and added to a sample to be analyzed in a 1:1 ratio (v/v) followed by heating for 5 min in a PCR cycler at 99 °C. Solution 3 was added in a 1:6 ratio (v/v), and the whole mixture was cooled down to room temperature before reading the absorbance at 575 nm on an Infinite M200 microplate reader (Tecan, Switzerland). The amount of reducing acids released is calculated based on a galacturonic acid standard curve. PG activity is expressed as nmol GalA equivalents/min/μg gut content protein released. Error bars indicate the standard error of the mean.

### RNA SEQ Analysis

RNA samples used for RNA-seq were the same as those used for RT-qPCR. Transcriptome sequencing was carried out for four biological replicates per treatment group and a total of 16 RNA samples using poly(A)+ enriched RNA fragmented to an average of 250 nucleotides. Sequencing was carried out by the Max Planck Genome Center, Cologne, on an Illumina HiSeq2500 Genome Analyzer platform using paired-end (2 x 150 bp) reads, yielding approximately 20-30 million reads for each of the 16 samples. Quality control measures, including the filtering of high-quality reads based on fastq file scores, the removal of reads containing primer/adapter sequences and trimming of the read length, were carried out using CLC Genomics Workbench v10.1 (https://www.qiagenbioinformatics.com/). The same software was used for *de novo* transcriptome assembly, combining randomly sampled batches of 15 Mio reads of two replicate samples each of the four RNA-seq treatment groups, using a total of 120 Mio reads and selecting the presumed optimal consensus transcriptome as previously described [42]. The final *de novo* reference gut transcriptome assembly (backbone) of *P.cochleariae* contained 72,572 contigs (minimum contig size = 250 bp) with an N50 contig size of 1,217 bp and a maximum contig length of 26,428 bp. The transcriptome was annotated using BLAST, Gene Ontology and InterProScan searches implemented in BLAST2GO PRO v5.1 (www.blast2go.de) as previously described [43].

Digital gene expression analysis was carried out using CLC Genomics workbench v10.1 to generate BAM mapping files, and QSeq (DNAStar Inc., US) to remap the Illumina reads from all 16 samples onto the reference transcriptome. The final step was to count the sequences to estimate the expression levels, using previously described parameters for mapping and normalization [44], but changing the read assignment quality options to require at least 80% of the total read bases and at least 90% of bases matching within each read to be assigned to a specific contig. To control for the effect of global normalization using the RPKM algorithm, we analyzed a number of highly conserved housekeeping genes, including those encoding GAPDH, ribosomal proteins Rps4e, Rps18 and Rpl7 and eukaryotic translation initiation factor 5a. The overall variation in expression level for these housekeeping genes was lower than 1.2-fold, indicating they were not differentially expressed. Rps18 and Rpl7 genes were used as reference gene and are shown in the heat map to confirm similar expression levels of control genes across treatment groups. The log2 (RPKM) values (normalized mapped read values; geometric means of the biological replicate samples) were subsequently used to calculate fold-change values. To identify differentially expressed genes, we used Student’s t-test (as implemented in Qseq) corrected for multiple testing using the Benjamini–Hochberg procedure to control the false discovery rate (FDR). The differential expression (fold-change values) of the GH28 genes, and the statistical significance thereof (Student’s t-test; FDR-corrected p-values), are summarized in Supplementary Table S2.

The short read data have been deposited in the EBI short read archive (SRA) with the following sample accession numbers: ERS2876704-ERS2876707. The complete study can also be accessed directly using the following URL: http://www.ebi.ac.uk/ena/data/view/PRJEB29501.

### Statistical analysis

The dependency of PG activity and GH28 expression levels on the different treatments was tested with one-way ANOVA analyses and the Tukey HSD test in order to find differences among the groups, both implemented in SigmaPlot 12.0. PG activity and expression level analyses are based on the means of technical replicates for each of the five biological replicates. Values for expression level of GH28-1 and GH28-4 were not normally distributed and failed the Equal Variance Test. The influence of different treatments on the expression of GH28-1 and GH28-4 were therefore investigated using the generalized least squares method (gls from the nlme library [45] to account for the variance in heterogeneity of the residuals. The varIdent variance structure (varIdent(form = ~1 | treatment)) was used. The influence of the treatment was determined by removing the explanatory variable and comparing the simpler model with the full model using a likelihood ratio test [46]. Differences between factor levels were determined by factor level reduction [47]. To compare weight gain over time in RNA*i*-treated larvae, we calculated the relative growth rate for a period of 5 days. The amount of leaf eaten was recorded at the same time. These two parameters were used to calculate the food-to-energy conversion efficiency. Statistical analyses were based on 50 replicates per treatment.

The dependency of developmental times (number of days until molting from 2nd to 3rd instar, termination of feeding, pupation, eclosion) on the treatment and the amount of consumed leaf material was determined by analyzing covariance with the different treatments as categorical and the amount of consumed leaf material as continuous explanatory variables. Differences between factor levels were determined by factor level reduction [47]. All data were analyzed with R version 3.4.1 [48].

## Results

### Silencing GH28 genes is specific

*P*. *cochleariae* has nine GH28 family members. Three act as endo-PGs (28-1, 5, 9), one as an oligogalacturonase (28-4) hydrolyzing trigalacturonic acid released by the endo-PGs (Kirsch et al., 2014). In addition, four GH28s (28-3, 6, 7, 8) do not show any activity towards pectic substrates or other PCW polysaccharides and possess amino acid substitutions in functionally important sites ([24], this study Fig. S1). To test for the biological impact of active and inactive GH28s, three genes each of active and inactive GH28 were silenced simultaneously: active endo-PGs, called 28a (*28-1, 5, 9*), and inactive GH28 pseudoenzymes, called 28i (*28-3, 6, 7*). Following RNAi, quantitative real-time PCR revealed a significant and specific down-regulation of target genes compared to the *GFP* injection control (Fig. 1A). It clearly illustrated the feasibility of simultaneously silencing several genes at once. In addition, the transcript abundances of GH28s not targeted through RNAi was not affected by any of the treatments, confirming the specificity of silencing genes of high sequence similarity (Fig. 1B).

**Fig. 1.**
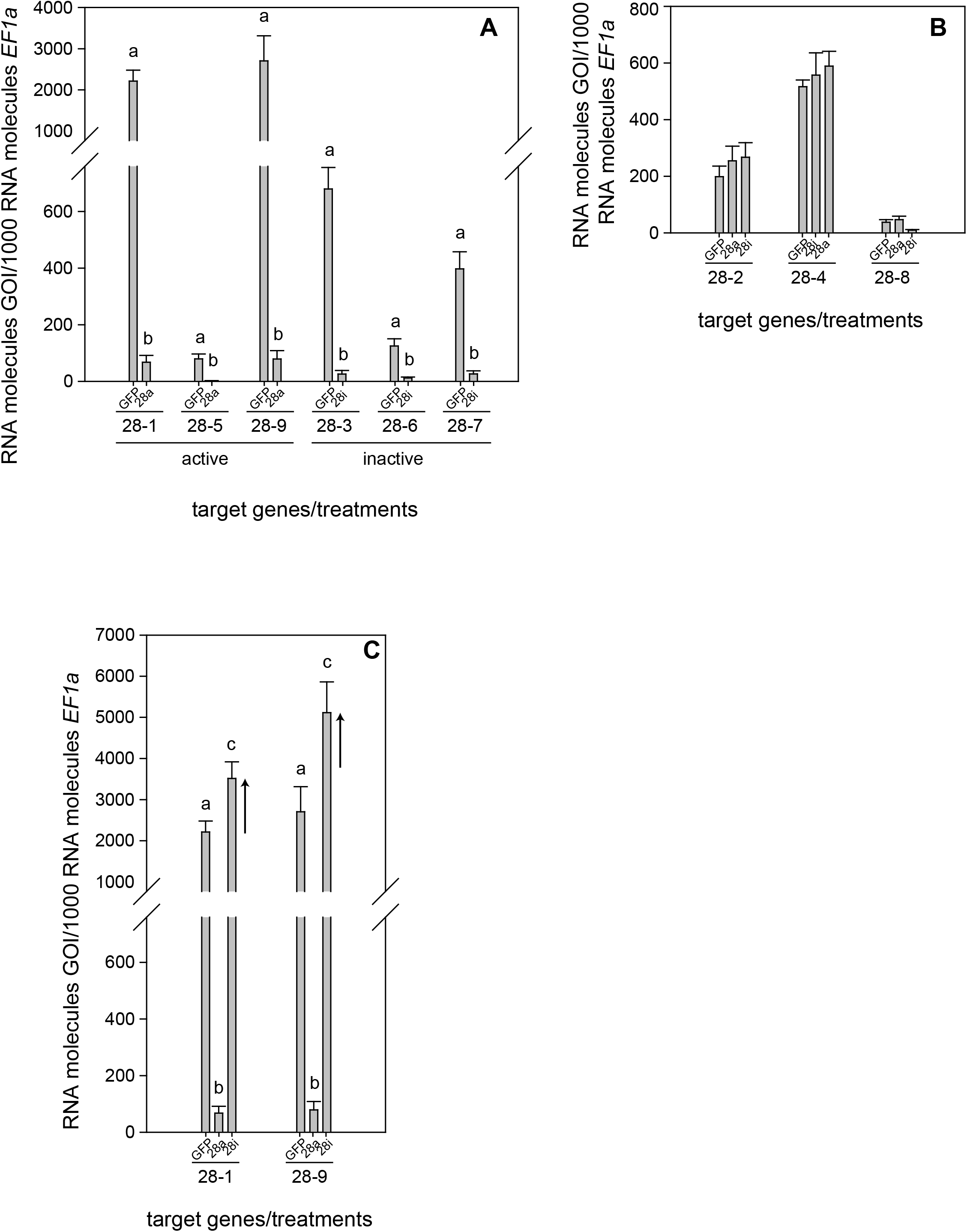
Expression pattern of *P*. *cochleariae* GH28s comparing injection control (GFP), active GH28 silencing (28a) and inactive GH28 silencing (28i). Expression of (A) GH28 targets, (B) untargeted GH28s and (C) the up-regulation of active GH28s when silencing their inactive GH28 counterparts. Transcript abundances are expressed as RNA molecules of gene of interest (GOI) per 1000 RNA molecules of the reference gene elongation factor 1-alpha (*EF-1α*).

### No global changes but treatment-specific GH28 induction

To obtain a more global view of silencing specificity as well as treatment-specific responses to gene expression levels, global gene expression analyses using RNA-Seq were performed. Transcript abundance was calculated based on four biological replicates for each of the treatments (GFP, 28a, 28i). We compared gene expression changes in 28a and 28i treatment samples relative to the GFP control but did not find a complex pattern of differentially expressed genes. More specifically, there were no significant gene expression changes except for the 28a and 28i targeted GH28 encoding genes. This gene expression pattern is exemplarily shown for transcripts encoding a variety of GH families (Fig. 2). Whereas targeted genes showed strong down-regulation compared to the control, the other GH families were not significantly affected. Among such families were further PCWDE-like cellulases (GH45) or xylanases (GH11), which obviously were not affected by the down-regulation of GH28s. Nevertheless, although not significant based on RNA-Seq data, the mRNA levels of the endo-PG 28-1 and 28-9 are higher in 28i than in the GFP control. To analyze this relationship in more detail, we performed qPCR to measure the expression of all GH28 genes for each treatment (Fig. S2). When the genes encoding active endo-PGs were silenced, the expression of the genes encoding the remaining GH28 family members (Fig. S2). Strikingly, when the genes encoding the inactive GH28s were down-regulated, two active endo-PGs (28-1, 9) were significantly up-regulated at the same time (Fig. 1C). Although 28-1 and 28-9 display the highest steady-state expression levels among all GH28s, they are still inducible to even higher levels (Fig. S2). In contrast, inactive GH28s are not differentially expressed when active PGs are silenced using RNAi. Thus, the differential expression of GH28s seems to depend on their function.

**Fig. 2.**
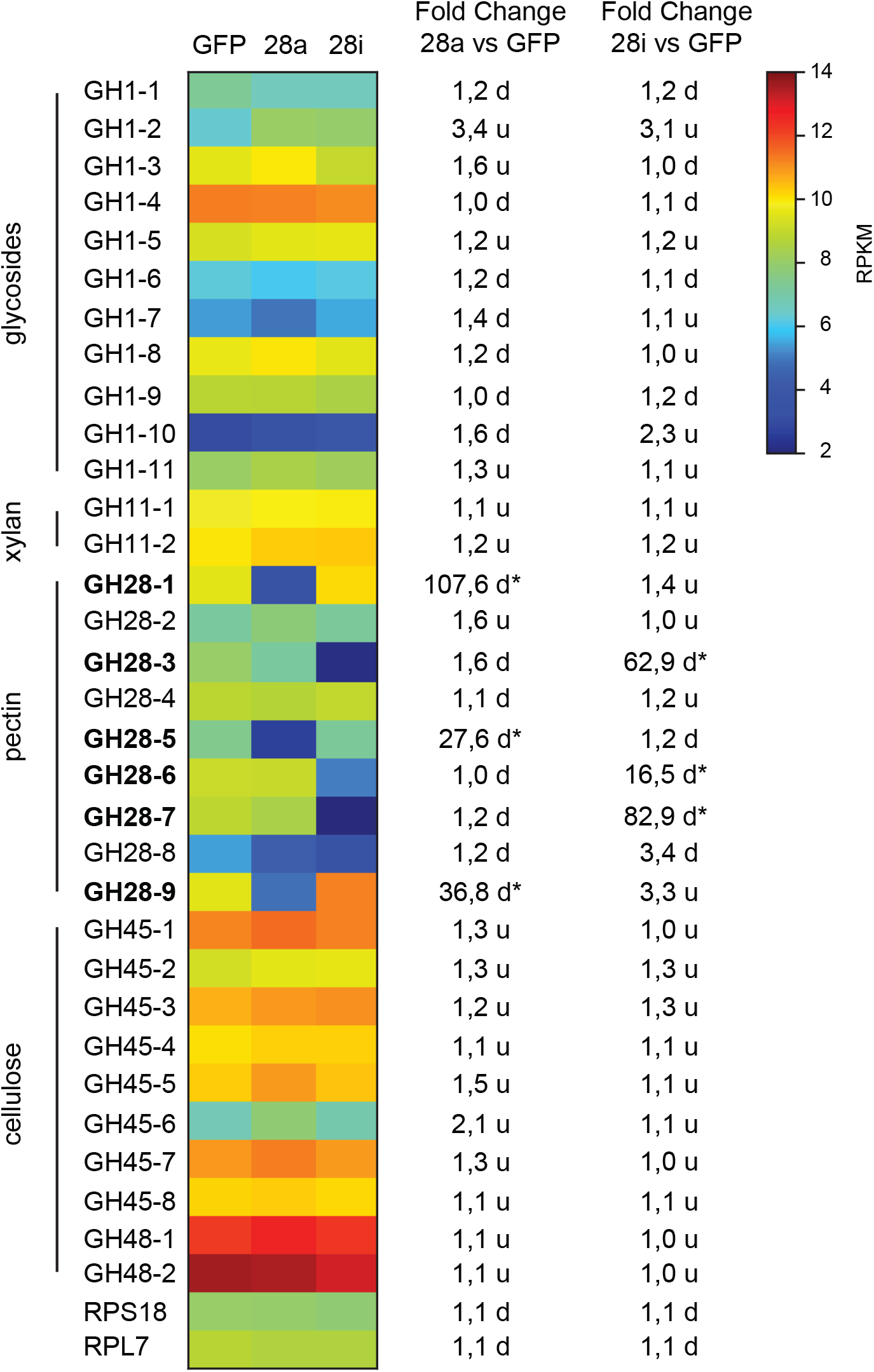
Heat map showing the relative expression levels of different glycoside hydrolase families (GH1 to 48) comparing the injection control (GFP) with the active (28a) and inactive (28i) GH28-silenced *P*. *cochleariae* larvae. Substrates of the different GH families are on the left. Silencing targets are indicated in bold and significant differences are shown with an asterisk (* < 0.05). Ribosomal protein 7 (RPL7) and ribosomal protein S18 (RPS18) are shown to confirm the uniform expression of these housekeeping genes across treatments. The map is based on log2-transformed RPKM values (blue represents weakly expressed genes, and red represents strongly expressed genes).

### Correlation of silencing with PG activity

To investigate the potential impact of gene silencing on gut PG activity, the release of polygalacturonic acid breakdown products by gut content was quantified. Endo-PG silenced larvae (28a) showed drastically reduced gut PG activity, with only about 10% remaining compared to the control (Fig. 3). These results confirmed that the recombinant proteins characterized previously as active endo-PGs (28-1, 5, 9) *in vitro* [24] were indeed responsible for the gut PG activity observed *in vivo*. PG activity correlated with the GH28 expression level in the 28a treatment. The PG activity in the 28i treatment did not differ from that of the GFP control. This similarity is surprising, as the up-regulation of the two dominant endo-PGs in the 28i treatment should have increased gut PG activity. Thus, PG activity seems not to correlate with the counter-regulation of the genes encoding the active PGs induced by the 28i silencing.

**Fig. 3.**
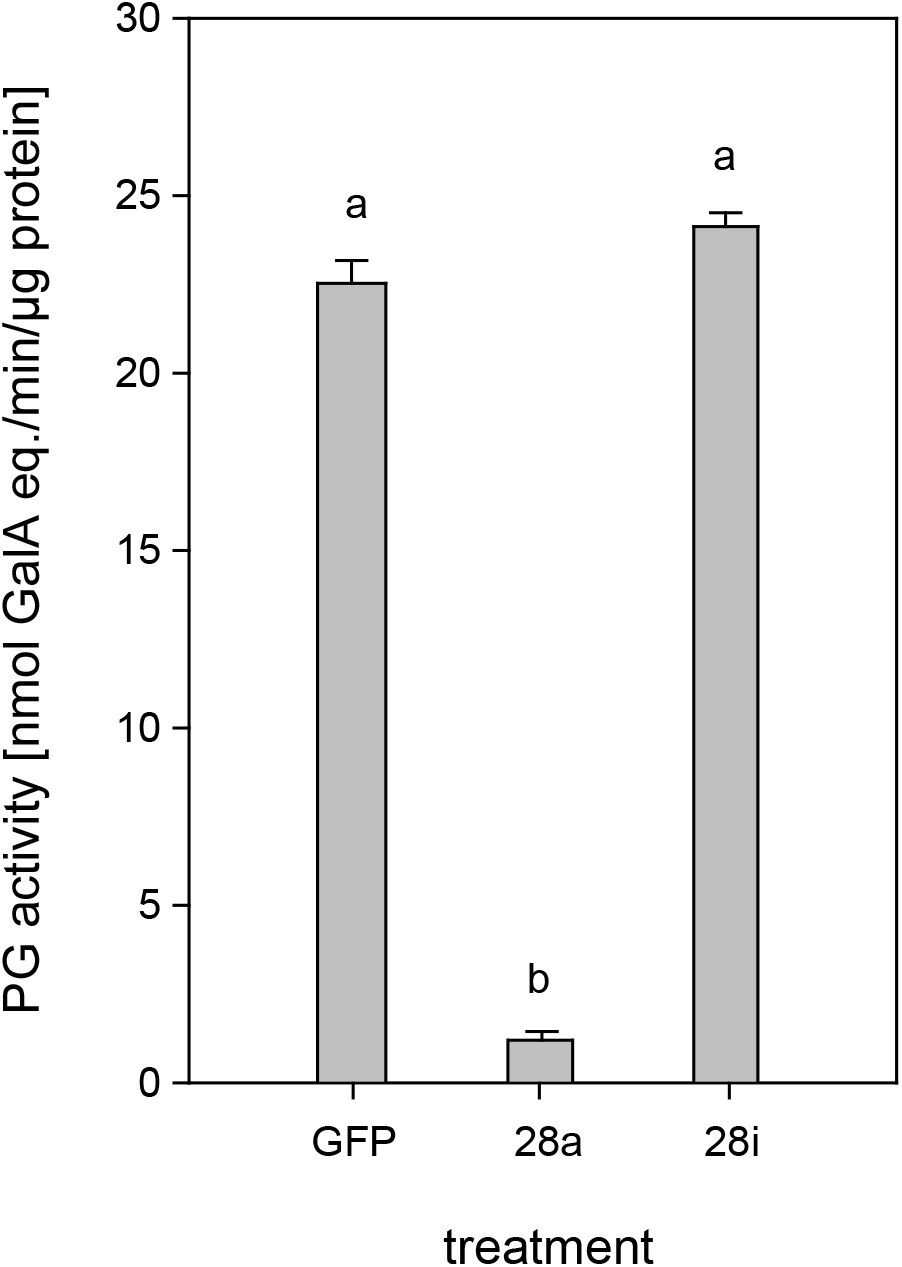
Quantification of PG activity in the *P*. *cochleariae* gut content. Hydrolytic activity in the larval guts of the injection control (GFP), active (28a) and inactive (28i) GH28-silenced larvae is shown. Activity is expressed in nmol-reducing uronic acids released per min and μg of gut content protein.

### Correlation of silencing with life history traits

The reduction of specific digestion-related enzymatic activity could lead to suboptimal nutrient release, which could in turn affect growth and development. To resolve the impact of impaired PG activity and thus less pectin breakdown in the gut, the amount of food ingested as well as the weight gain of *P*. *cochleariae* larvae were recorded. The change in weight as a function of the amount of food ingested is an indicator of food-to-energy conversion efficiency and thus a measurement of how efficiently nutrients can be released in the gut and subsequently used. The weight gain per cm^2^ leaf eaten was significantly lower in the 28i treatment, compared with the 28a treatment (Fig. 4), indicating a less efficient food-to-energy conversion in the larvae for which the inactive PGs were silenced. Although inactive GH28s presumably do not impact pectin hydrolysis, our results suggest they have an important function in processing plant material and digestion. This connection seems counter-intuitive, as the silencing of pseudoenzymes seems to have a higher impact on food-to-energy conversion efficiency than the silencing of their active relatives. To test if these differences among treatments influenced larval development, we recorded time to pupation and time to eclosion in days after experimental injections. We further tested if developmental time depends on the treatment and the food consumed by the analysis of covariance. The time until pupation and eclosion depended on the treatment (pupation: F=24.058, p<0.001; eclosion: F= 5.692, p=0.001) and the consumed food (pupation: F= 6.673, p<0.001; eclosion: F= 14.538, p<0.001) (Fig. 5A, B). More precisely, the larvae for which inactive GH28s were silenced developed more slowly with the same amount of food ingested compared to the other treatments, none of which showed any difference. Thus, the silencing of inactive GH28s prolongs the developmental period, which supports the important function of inactive GH28s.

**Fig. 4.**
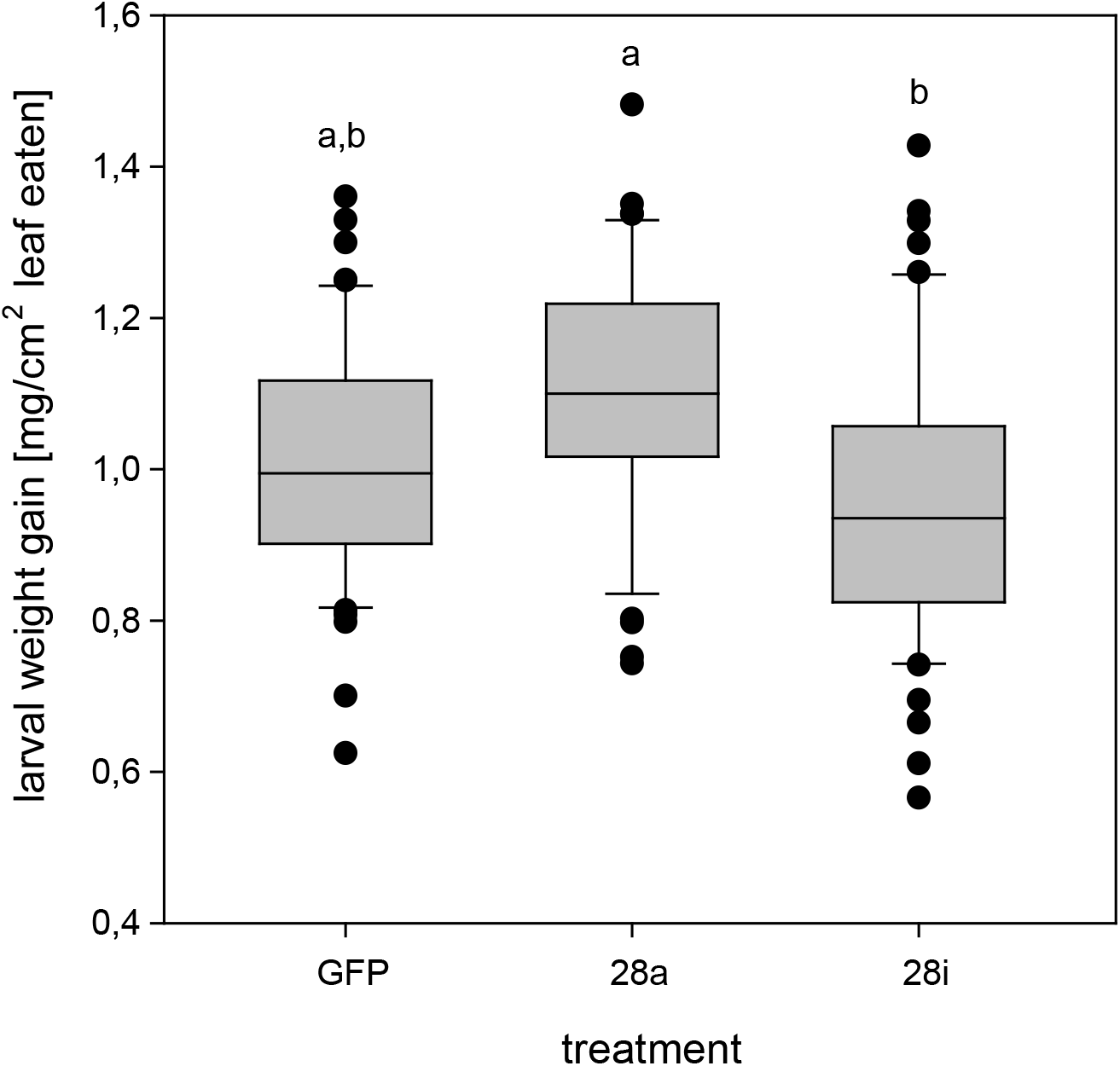
Efficiency of food-to-energy conversion from early 2nd- to 3rd-instar *P*. *cochleariae* larvae. Injection control (GFP) and the two silencing treatments are compared and the efficiency is calculated as mg larval weight gain per cm^2^ leaf eaten over time.

**Fig. 5.**
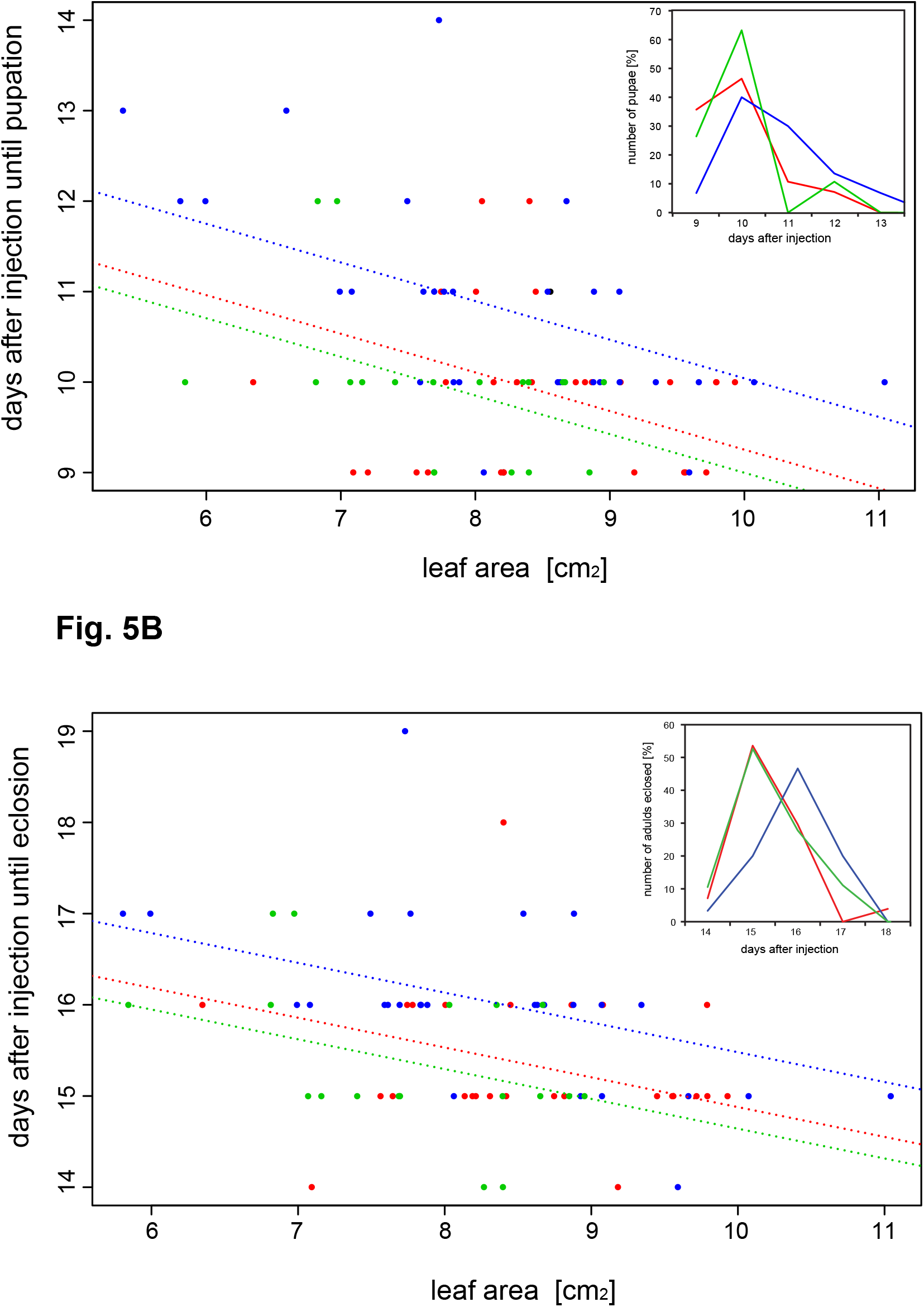
Dependency of (A) time to pupation and (B) time to eclosion from the amount of the consumed leaf area and the treatment (GFP: red, 28a: green and 28i: blue). Insertions show the number of pupae and adults emerged in percentages over days after injection, respectively.

## Discussion

Herbivorous insects have an optimal carbon-to-nitrogen (C/N) ratio for food intake, which they often fail to achieve due to a unbalanced C/N plant diet [6]. In plants, starch is the main storage polysaccharide and the major source of carbohydrate-based energy in herbivorous insects. Fitness costs, such as developmental delay resulting from altered starch digestion in insects either through the ingestion of plant α-amylase inhibitors or the knockdown of amylase genes, illustrate the importance of carbohydrate accessibility [49-51].

There is another potential carbohydrate source that is omnipresent in a herbivore’s diet but often overlooked: the plant cell wall (PCW). The PCW is rich in polysaccharides such as cellulose, various hemicelluloses and pectin, and is highly conserved [52]. PCW-degrading enzymes (PCWDEs) are widely distributed in insects [12]. Since many insects rely on a nitrogen-poor diet that is rich in cellulose, it is conceivable that herbivorous insects exploit this source of carbohydrates [53]. Experiments with feeding termites a 13C-labelled cellulose diet showed that the 13C labels appear fixed in amino acids supplemented by the termites’ gut microbes, providing evidence that cellulose degradation increases nitrogen levels and is beneficial for some insects [54]. In addition, silencing of a cellulase in larvae of the western corn rootworm *Diabrotica v. virgifera* lowered weight gain and increased the time to pupation compared to control larvae [55]. These results have to be taken with caution as the authors neither run off-target predictions nor showed a reduction of cellulase activity resulting from gene silencing. Thus, the observed phenotype cannot be correlated with lower cellulase activity with total certainty. Moreover, the knockdown of GH45 cellulases in the leaf beetle *Gastrophysa viridula* had no effect on larval fitness [56],), indicating that the impact of cellulase activity in herbivorous insects depends on both species and context, such as the diet provided for feeding assays..

In addition to cellulose, herbivorous insects feeding on living plant material ingest, high amounts of pectin. Whether insects benefit from pectin digestion is not clear. We found no effect on *P. cochleariae* larval development when silencing active PGs in the 28a treatment. Surprisingly, we detected an effect on insect fitness and on gene expression when silencing the inactive GH28s compared to the active ones. Furthermore, silencing GH28 pseudoenzymes lowered the food-to-energy conversion efficiency and lengthened the time required for development. These effects are similar to biological consequences caused by the suppression of digestive enzymes such as the amylases mentioned above, as well as alpha-glucosidases [57], proteases [58], and lipases [59]. At first glance, the presence of fitness costs in insects of the 28i but not in those of the 28a treatment is counter-intuitive, as the silencing efficiency is comparable between the two treatments and the suppression of the PGs goes hand in hand with a drastic reduction in PG activity.

Gut enzymes are usually part of an intertwined and finely tuned digestive system, which can be regulated at multiple levels and in a manner that is not predictable. For example, the inhibition of proteases of the Phytophaga seed beetle *Callosobruchus maculatus* resulted in the differential expression of PCWDEs, including GH5 mannanases and GH28s [60, 61]. This effect on PCWDE expression indicates crosstalk between digestive enzymes involved in protein and polysaccharide breakdown, and, even more importantly, illustrates the impact of PCWDEs in insects coping with sub-optimal diets. The performance of *P. cochleariae* depends on host plant species as well as on plant quality [62, 63] and pectin amount and structure generally differ between plants [64, 65]. The *P. cochleariae* laboratory strain used in our experiments is kept under optimal conditions and is adapted to *Brassica* species used for rearing since many generations. Therefore, decreased PG activity might have a strong effect on the larvae exposed to a challenging diet and ecologically relevant environment. The movement of the food bolus, and with that the amount of time PGs and pectin can interact in the gut, is highly variable in insects, ranging from hours to several days [3]. Thus, although PG activity in the 28a treatment is impaired, the interaction time of PGs and pectin in the gut might be enough for sufficient enzyme function. As “Phytophaga” beetles also possess gene families encoding active cellulases and hemicellulases [25, 56, 66-69], it is unclear whether the lack of a specific PCWDE activity, such as that of PGs, is costly or can be compensated for by the concurrent action of other enzymatic functions.

As the function of the inactive GH28s is unknown, the diminished food-to-energy conversion efficiency and extended developmental delay caused by their silencing is hard to explain. Although challenging [1], the functions already assigned to pseudoenzymes are extraordinarily diverse, ranging from regulators of their active counterparts or inhibitors of completely unrelated enzymes to being decoys that snatch away inhibitors to protect closely related enzymes [70-72]. To clarify the role of the inactive GH28 pseudoenzymes, we performed RNA-Seq-based global gene expression analysis, combined with qRT-PCR analyses of selected genes, to identify treatment-dependent changes in gene expression. Surprisingly, when treatments with the gfp injection control were compared, we did not find a complex pattern of differentially expressed genes in our analysis of the global transcriptome data. The only significant changes in gene expression are found in the GH28 genes that were down-regulated in the corresponding treatments. In addition, we found a slight up-regulation of active PGs when knocking down the inactive GH28s. qPCR analyses revealed that the two active PGs (28-1 and 28-9) are significantly up-regulated in the 28i treatment. This induction indicates a crosstalk between active and inactive GH28 family members and shows that the expression levels depend on each other at least in one direction. However, the up-regulation of the expression of PGs in the 28i treatment does not fit with the observed PG enzyme activity levels. Although PG activity should be higher in the 28i compared to the gfp control treatment, due to higher PG expression levels, the activity levels do not differ. The only plausible explanation for this discrepancy is that the down-regulation of the inactive GH28s results in a reduction of PG activity in the gut, which is compensated for by the up-regulation of PGs. The possibility thus exists that the inactive GH28 proteins, although pseudoenzymes, are still linked to the pectolytic pathway, which, at least in part, could explain the observed developmental delay. Additional support for the synergistic character of active and inactive GH28s comes from the temporal and spatial co-expression of those genes [24, 28, 30].

Our data reveal that the loss of enzymatic activity towards an ancestral substrate does not mean that these proteins have completely lost their impact on the underlying pathway but, rather, suggests they may be important for the organism. The high steady-state expression level of active endo-PGs and their dynamic regulation stand for the robustness of the pectolytic system and thus its importance for herbivorous beetles. Knocking down inactive GH28s exceeds the impact of their active counterparts, which is a second layer of insurance to encourage pectin digestion.

---

**Fig. S1.**
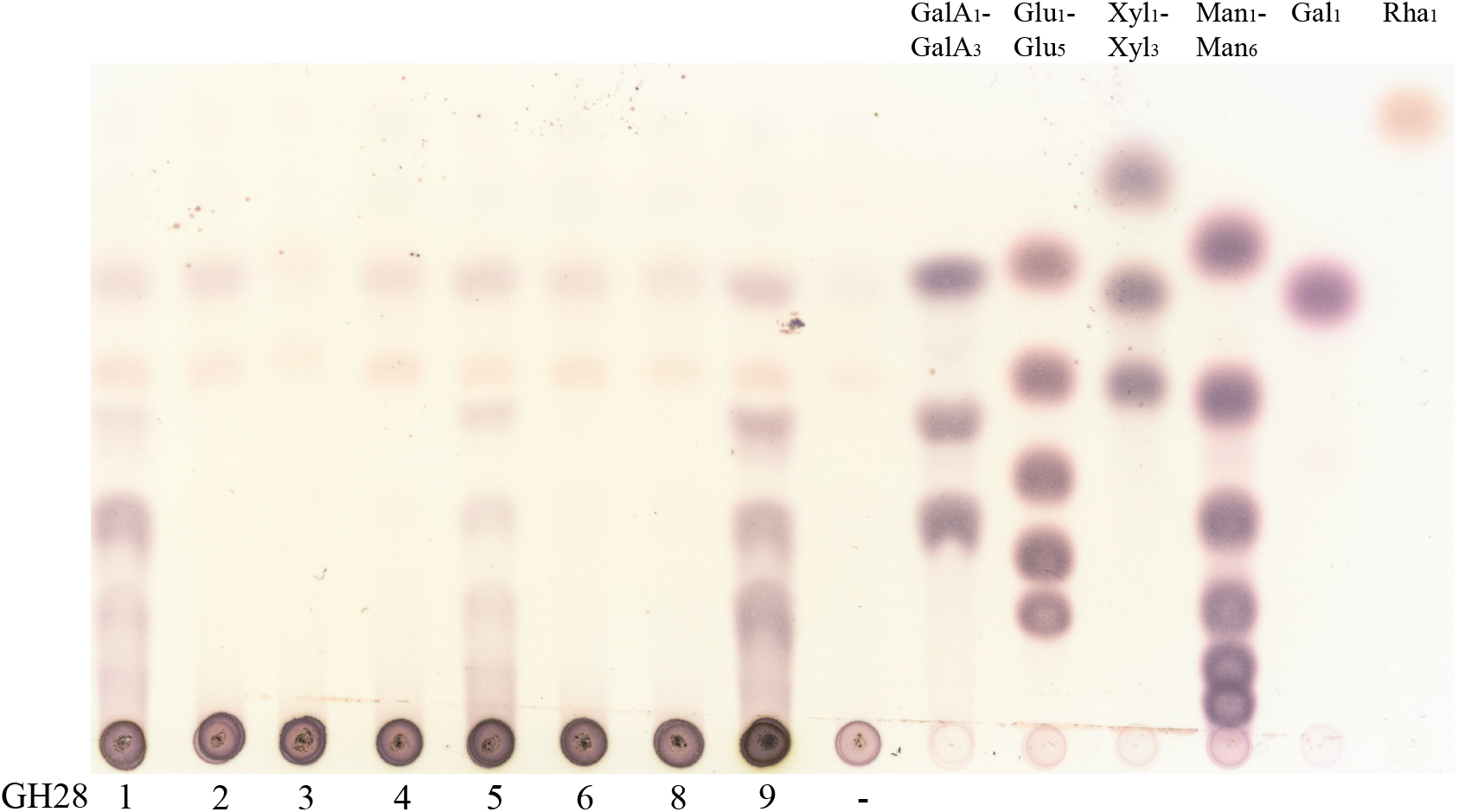
Activity of *P. cochleariae* GH28s onthe Chinese cabbage PCW substrate. Analysis of breakdown products of GH28s by thin-layer-chromatography (TLC) is shown. Standards of breakdown products of the following substrates were used: pectic polygalacturonan (GalA1-3), cellulose (Glu1-5), xylan (Xyl1-3), mannan (man1-6) as well as galactose (Gal) and rhamnose (Rha) monomers, which are decorations of pectin.

**Fig. S2.**
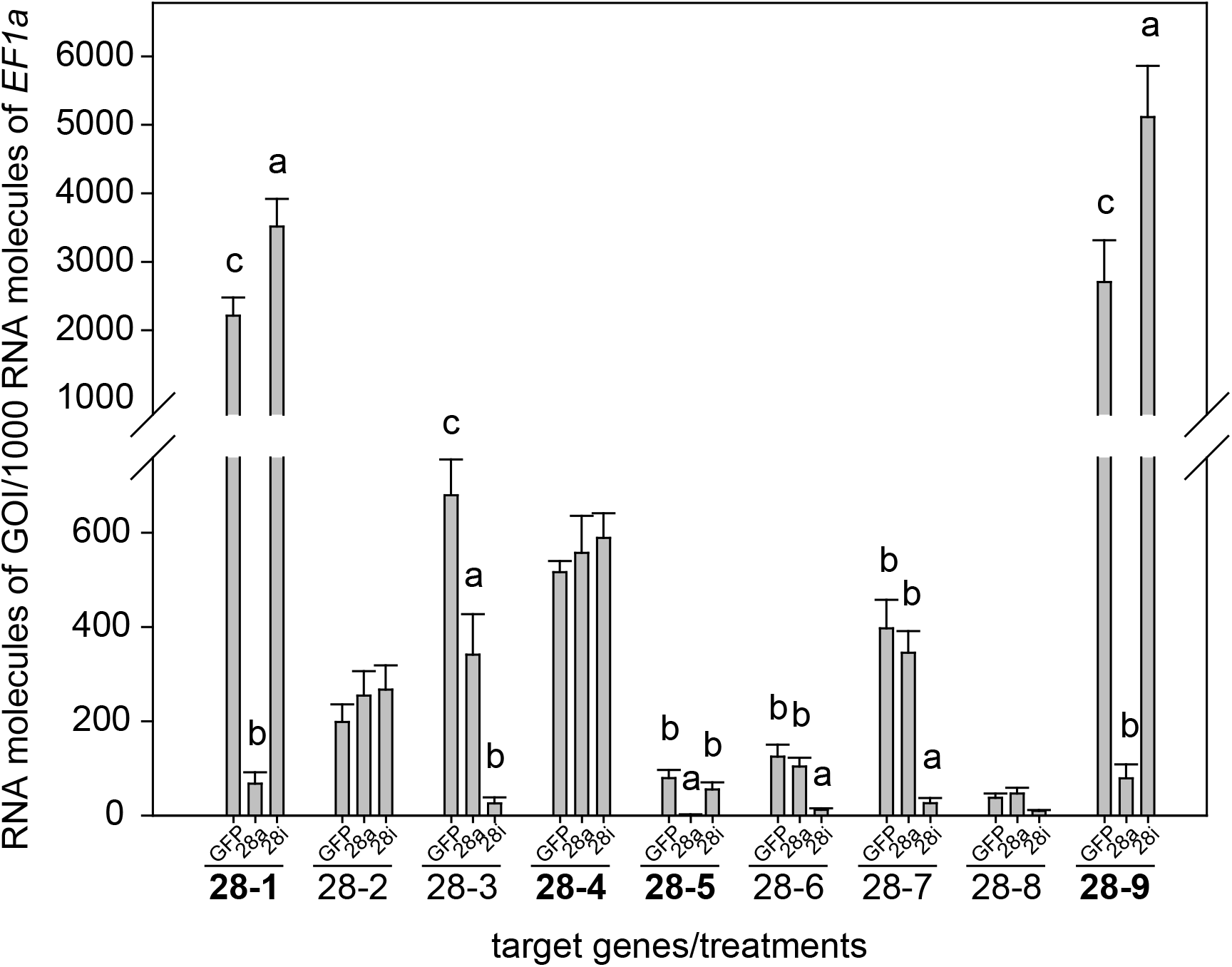
General overview of the expression patterns of all *P*. *cochleariae* GH28s over all treatments comparing injection control (GFP), active GH28 silencing (28a) and inactive GH28 silencing (28i). Active GH28s are indicated in bold. Transcript abundances are expressed as RNA molecules of gene of interest (GOI) per 1000 RNA molecules of the reference gene elongation factor 1-alpha (*EF-1α*).

**Table S1.**
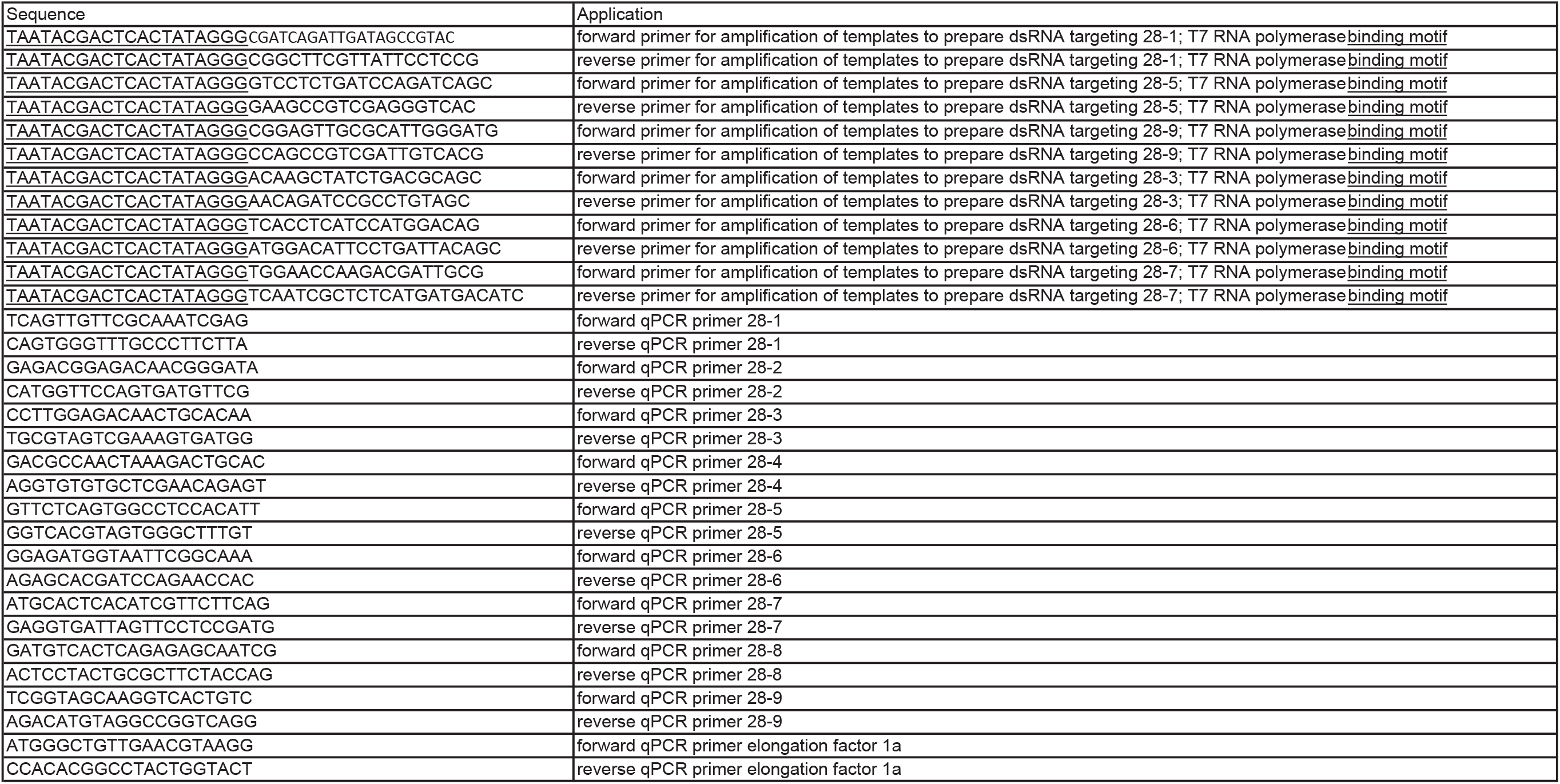
List of primers used for dsRNA synthesis and qRT-PCR of *P*. *cochleariae* GH28s.

**Table S2.**

| description | Pco-ActiveGH28 vs PCO-GFP - FoldChange | Pco-ActiveGH28 vs PCO-GFP -P value |
| --- | --- | --- |
| GH1-1 | 1,172 down | 1 |
| GH1-2 | 3,361 up | 0.593 |
| GH1-3 | 1,578 up | 1 |
| GH1-4 | 1,031 down | 1 |
| GH1-5 | 1,164 up | 0.992 |
| GH1-6 | 1,152 down | 0.909 |
| GH1-7 | 1,386 down | 1 |
| GH1-8 | 1,228 up | 1 |
| GH1-9 | 1,020 down | 1 |
| GH1-10 | 1,561 up | 1 |
| GH1-11 | 1,284 up | 0.909 |
| GH11-1 | 1,073 up | 1 |
| GH11-2 | 1,214 up | 1 |
| GH28-1 | 107,550 down | 0.0256 |
| GH28-2 | 1,555 up | 0.968 |
| GH28-3 | 1,557 down | 1 |
| GH28-4 | 1,083 down | 1 |
| GH28-5 | 27,618 down | 0.0257 |
| GH28-6 | 1,022 down | 1 |
| GH28-7 | 1,215 down | 1 |
| GH28-8 | 1,233 down | 1 |
| GH28-9 | 36,759 down | 0.0196 |
| GH45-1 | 1,266 up | 1 |
| GH45-2 | 1,255 up | 1 |
| GH45-3 | 1,245 up | 1 |
| GH45-4 | 1,110 up | 1 |
| GH45-5 | 1,527 up | 1 |
| GH45-6 | 2,143 up | 1 |
| GH45-7 | 1,268 up | 1 |
| GH45-8 | 1,058 up | 1 |
| GH48-1 | 1,114 up | 1 |
| GH48-2 | 1,100 up | 1 |
| RPS18 | 1.059 down | 1 |
| RPL7 | 1.114 down | 1 |

| description | Pco-InactiveGH28 vs PCO-GFP - FoldChange | Pco-InactiveGH28 vs PCO-GFP - P value |
| --- | --- | --- |
| GH1-1 | 1,178 down | 1 |
| GH1-2 | 3,114 up | 0.393 |
| GH1-3 | 1,015 down | 0.893 |
| GH1-4 | 1,132 down | 0.893 |
| GH1-5 | 1,216 up | 1 |
| GH1-6 | 1,078 down | 1 |
| GH1-7 | 1,098 up | 1 |
| GH1-8 | 1,029 up | 0.924 |
| GH1-9 | 1,235 down | 1 |
| GH1-10 | 2,311 up | 1 |
| GH1-11 | 1,060 up | 1 |
| GH11-1 | 1,097 up | 1 |
| GH11-2 | 1,232 up | 1 |
| GH28-1 | 1,426 up | 0.899 |
| GH28-2 | 1,035 up | 1 |
| GH28-3 | 62,886 down | 0.0226 |
| GH28-4 | 1,122 up | 0.987 |
| GH28-5 | 1,151 down | 1 |
| GH28-6 | 16,509 down | 0.0247 |
| GH28-7 | 82,876 down | 0.0118 |
| GH28-8 | 3,441 down | 1 |
| GH28-9 | 3,305 up | 1 |
| GH45-1 | 1,030 up | 1 |
| GH45-2 | 1,300 up | 1 |
| GH45-3 | 1,328 up | 1 |
| GH45-4 | 1,142 up | 1 |
| GH45-5 | 1,072 up | 1 |
| GH45-6 | 1,115 up | 1 |
| GH45-7 | 1,027 down | 1 |
| GH45-8 | 1,057 down | 1 |
| GH48-1 | 1,026 up | 1 |
| GH48-2 | 1,005 up | 1 |
| RPS18 | 1.137 down | 1 |
| RPL7 | 1.117 down | 1 |

## Author contributions

R.K. conceived and designed the experiments; R.K. and H.V. performed the experiments; R.K., H.V. and G.K. analyzed the data; R.K. wrote the paper with major contributions from all co-authors. All authors read and approved the final manuscript.

## Acknowledgments

We thank Bianca Wurlitzer for *P*. *cochleariae* stock rearing and the greenhouse team for plant rearing. We thank Caroline Müller, David Heckel and Franziska Beran for fruitful discussions regarding experimental setup and Wiebke Häger for preparing the PCW substrate. We thank Emily Wheeler for editorial assistance. Financial support from the Max Planck Society and the German Science Foundation (KI1917/1-1) is gratefully acknowledged.

